# Ovarian status dictates the neuroinflammatory and behavioral consequences of sub-chronic stress exposure in middle-aged female mice

**DOI:** 10.1101/706887

**Authors:** Rand S. Eid, Stephanie E. Lieblich, Sarah J. Wong, Liisa A.M. Galea

## Abstract

Ovarian hormones influence the outcomes of stress exposure and are implicated in stress-related disorders including depression, yet their roles are often complex and seemingly contradictory. Importantly, depression and stress exposure are associated with immune dysregulation, and ovarian hormones have immunomodulatory properties. However, how ovarian hormones can influence the inflammatory outcomes of stress exposure is poorly understood. Here, we examined the effects of long-term ovariectomy on the behavioral and neuroinflammatory outcomes of sub-chronic stress exposure in middle-aged mice. Briefly, sham-operated and ovariectomized mice were assigned to non-stress groups or exposed to 6 days of variable stress. Mice were assessed on a battery of behavioral tests, and cytokine concentrations were quantified in the frontal cortex and hippocampus. In the frontal cortex, postsynaptic density protein-95 expression was examined as an index of excitatory synapse number and/or stability, and phosphorylated mitogen-activated protein kinases (MAPKs) were measured to explore potential cell signaling pathways elicited by stress exposure and/or ovarian hormones. Long-term ovariectomy modified the central cytokine profile by robustly reducing cytokine concentrations in the frontal cortex and modestly increasing concentrations in the hippocampus. Under non-stress conditions, long-term ovariectomy also reduced extracellular signal-regulated kinase (ERK) phosphoprotein expression in the frontal cortex and increased some measures of depressive-like behavior. The effects of sub-chronic stress exposure were however more pronounced in sham-operated mice. Notably, in sham-operated mice only, sub-chronic stress exposure increased IL-1β and IL-6:IL-10 ratio in the frontal cortex and hippocampus and reduced pERK1/2 expression in the frontal cortex. Further, although sub-chronic stress exposure increased anhedonia-like behavior regardless of ovarian status, it increased passive-coping behavior in sham-operated mice only. These data indicate that long-term ovariectomy has potent effects on the central cytokine milieu and dictates the neuroinflammatory and behavioral effects of sub-chronic stress exposure in middle-aged mice. These findings therefore suggest that the immunomodulatory properties of ovarian hormones are of relevance in the context of stress and possibly depression.

## Introduction

Major Depressive Disorder (MDD), a leading cause of disability and ill-health worldwide (Friedrich, 2017), is approximately twice more prevalent in women than men (Kessler and Bromet, 2013; Salk et al., 2017). In addition to this robust difference in prevalence rates, disease presentation and pathophysiology differ significantly between men and women (reviewed in Eid et al., 2019). Although there is an encouraging increase in studies investigating sex differences in depression and in animal models of stress exposure, studies that target female-specific factors that may contribute to risk or resilience to depression remain scarce. Ovarian hormones play a role in mood regulation and influence depression in women (Brummelte and Galea, 2010; Hantsoo and Epperson, 2015; Soares, 2013) and therefore represent one such target. For example, there is an increased susceptibility to develop depression during the reproductive years in women compared to men (Gutiérrez-Lobos et al., 2002). These data provide the basis for the hypothesis that ovarian hormones may increase risk for depression in women. Other lines of evidence, however, support the contrasting hypothesis that ovarian hormones may afford resilience against depression. Specifically, events that are characterized by large reductions in ovarian hormones, including the postpartum and perimenopause, are associated with an increased risk for depression in women (Cohen et al., 2006; Freeman et al., 2004; Hendrick et al., 1998; Soares, 2014). In rodent studies, ovariectomy alone increases depressive-like behavior in the short term and increases susceptibility to the development of depressive-like behavior in response to chronic stress exposure in the long term (Lagunas et al., 2010; Li et al., 2014; Mahmoud et al., 2016). Taken together, this literature highlights the complex and seemingly paradoxical roles of ovarian hormones in mood regulation and depression. Importantly, ovarian hormones regulate a wide range of physiological processes and systems that are compromised in depression, including stress response and immune systems (Goel et al., 2014; Klein and Flanagan, 2016). Therefore, examining the interplay between endocrine and immune systems in the context of stress exposure may clarify the role of ovarian hormones in depression.

There is mounting evidence of immune dysregulation in depression (Hodes et al., 2015; Miller and Raison, 2016). At least a subpopulation of individuals with MDD present with increased markers of inflammation, supported by several meta-analyses which highlight that IL-6 and TNF-α are increased in the blood and cerebral spinal fluid (Dowlati et al., 2010; Haapakoski et al., 2015; Liu et al., 2012; Wang and Miller, 2018). Further, poor antidepressant efficacy has been linked to an inability to normalize dysregulated inflammatory processes (Syed et al., 2018). Stress exposure, a major risk factor for depression (Kendler et al., 1999), can also activate inflammatory responses in the brain and periphery (Felger et al., 2015; Gouin et al., 2012). Importantly, sex differences exists across many immune processes, mediated in part by sex hormones (Klein and Flanagan, 2016). Several reports document the anti-inflammatory actions of estrogens and progesterone in the brain; in response to acute inflammatory challenges *in vitro* and *in vivo*, estradiol, selective estrogen receptor modulators (SERMs), and progesterone have all been shown to ameliorate microglia inflammatory activity and the secretion of proinflammatory cytokines including IL-1β, TNF-α, and IL-6 (Lei et al., 2014; Smith et al., 2011; Suuronen et al., 2005; Tapia-Gonzalez et al., 2008; Vegeto et al., 2001). Further, in models of ischemic stroke and brain injury, both estrogens and progesterone have well-documented neuroprotective and anti-inflammatory properties (Sayeed and Stein, 2009; Villa et al., 2016). However, the immunomodulatory actions of estrogens and progesterone are complex, as they can produce pro-inflammatory and neurotoxic effects depending on many factors, including age and sex (Hsieh et al., 2016; Selvamani and Sohrabji, 2010a; Straub, 2007). Despite the recognition that ovarian hormones influence immune processes, the role of ovarian hormones at the intersection of depression and immune dysregulation is not well known. This is particularly important during the transition to menopause, a time of heightened risk for MDD (Cohen et al., 2006; Freeman et al., 2004) that is also marked by substantial changes in ovarian and immune function (Burger et al., 2002; Giefing-Kröll et al., 2015). The current study targets this gap in the literature by examining how the ovarian hormone milieu can influence the depressogenic and inflammatory consequences of stress exposure in middle-aged mice.

A better understanding of endocrine-immune interactions in the pathogenesis of depression may be achieved by investigating downstream intracellular signaling pathways that are elicited by stress exposure. Mitogen-activated protein kinase (MAPK) pathways represent a logical target, as they regulate a wide range of cellular processes in response to diverse stimuli, including hormones and cytokines (Cargnello and Roux, 2011). Further, studies in humans and animal models of depression indicate that MAPK signaling pathways are compromised in the disorder (Hollos et al., 2018; Miller and Raison, 2006; Wang and Mao, 2019). The extracellular signal-regulated kinase (ERK) subfamily of MAPKs has received most attention in the context of depression (Wang and Mao, 2019), however the stress-activated protein kinases, c-Jun NH_2_-terminal kinase (JNK) and p38 MAPK, may also be implicated (Hollos et al., 2018; Miller and Raison, 2006). Here, we quantified phosphorylated MAPKs to explore cellular mechanisms that may underly the role of ovarian hormones in risk or resilience in the face of stress exposure.

Disruptions in synaptic function and number are evidenced following stress and thought to be key factors in the pathogenesis of stress-related disorders such as depression (Duman et al., 2016). The prefrontal cortex (PFC) may be particularly vulnerable; in addition to reports of reduced PFC volume in depression (Drevets, 2000; Drevets et al., 2008), post-mortem studies find reductions in the number of synapses and in the expression of synapse-related genes in the PFC (Feyissa et al., 2009; Kang et al., 2012). In rodent models, chronic stress exposure results in dendritic remodeling in the frontal cortex and hippocampus (reviewed in McEwen et al., 2016), in addition to alterations is spines and synaptic markers (Orlowski et al., 2012; Workman et al., 2013). Importantly, neuronal remodeling in the frontal cortex and hippocampus in response to stress is influenced by sex and sex hormones (Galea et al., 1997; Shansky et al., 2010, 2009). Further, inflammatory processes could be implicated in synaptic alterations in stress and depression, as synaptic pruning is mediated by microglia during development and possibly in disease (Hong et al., 2016; Paolicelli et al., 2011). Therefore, examining synaptic markers may also clarify endocrine-immune interactions in stress and depression.

In this study, we used female mice to examine the behavioral and inflammatory consequences of sub-chronic stress exposure in middle age, a time when intact animals transition into reproductive senescence. We chose middle age because few studies have examined this important time point in females, and because perimenopause represents a time of increased vulnerability to depression in women. We measured cytokine concentration in the periphery, the hippocampus, and frontal cortex, areas known to be susceptible to stress exposure and compromised in depression (Howard et al., 2019; McKinnon et al., 2009). We also quantified phosphorylated MAPKs and postsynaptic density protein 95 (PSD-95) in the frontal cortex. We hypothesized that ovarian hormone deprivation will dictate behavioral outcomes and modulate cytokines, cell signaling proteins, and PSD-95 expression in response to stress exposure.

## Method

### Animals and surgery

Young-adult female C57BL/6N mice were purchased from Charles River Laboratories (Quebec, Canada). Mice received bilateral ovariectomy (OVX; n = 14) or sham surgery (Sham; n = 14) at Charles River Laboratories in young adulthood at 8 weeks of age, and arrived at our facility at 9 weeks of age. Mice were group housed (2-3/cage) in a temperature- and humidity-controlled colony room (21 ± 1°C; 50 ± 10% humidity), maintained on a 12-hour light/dark cycle (lights on at 07:00 h), and provided *ad libitum* access to food and water. Mice were left undisturbed, apart from weekly cage changing, until the beginning of further experimental manipulations in middle age, at 11-months-old. The reason mice were tested at middle age was two fold: 1) this is a time of heightened vulnerability to depression in women (Cohen et al., 2006; Freeman et al., 2004) that few animal studies target, and 2) previous rodent studies suggest that differences in susceptibility to stress exposure may emerge only after longer periods of ovarian hormone deprivation (Lagunas et al., 2010). Although our ovariectomized groups may not closely model the menopausal transition as surgery was performed in young adulthood, our sham-operated groups were tested during the transition to reproductive senescence, and as such are an appropriate model for the menopausal transition (Koebele and Bimonte-Nelson, 2016). All procedures were approved by the Animal Care Committee at the University of British Columbia and were performed in accordance with the ethical guidelines set by the Canadian Council on Animal Care.

### Sub-chronic variable stress exposure

In each ovarian status condition (OVX, Sham), half the mice were exposed to 6 days of sub-chronic variable stress, adapted from previous methods (Hodes et al., 2015; Labonté et al., 2017). We chose a 6-day paradigm to examine group differences in early indicators of immune dysregulation in response to stress. Mice in the stress condition were transferred to a separate colony room and allowed to acclimatize for two weeks prior to stress exposure. Stress-exposed mice were housed separately to avoid the transmission of stress-induced olfactory cues to non-stressed mice (Brechbuhl et al., 2013). The protocol consisted of three stressors, applied once per day on days 1-3, and repeated on days 4-6 (see timeline in **Fig. 1**). Mice were exposed to foot shock stress on days 1 and 4 (100 shocks at 0.45mA, randomly distributed across 1 hour), tail suspension stress on days 2 and 5 (1 hour, suspended using laboratory tape, mid-way across the tail), and restraint stress on days 3 and 6 (1 hour, in a well-ventilated 50ml falcon tube, in the home cage). Mice in non-stress groups were handled daily for 5 minutes across days 1-6, but otherwise left undisturbed. Body weight was monitored daily across the stress exposure period and in non-stressed controls.

**Figure 1.**
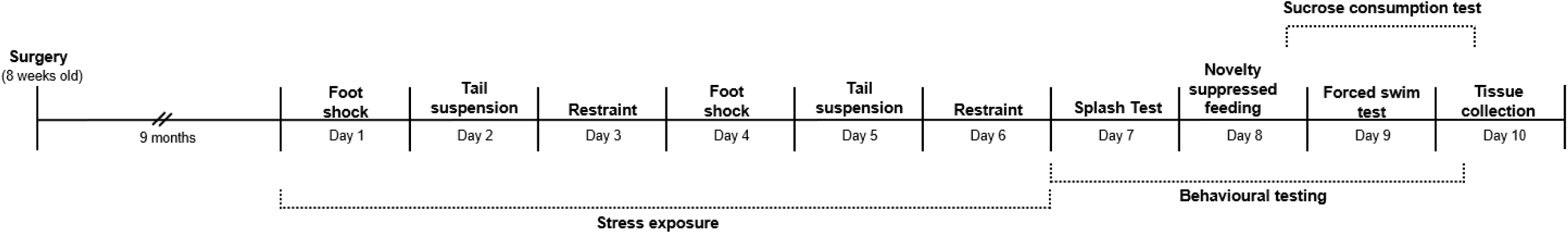
Experimental timeline. Mice received bilateral ovariectomy or sham surgery in young adulthood (8-weeks-old). In middle age (11-months-old), mice in the stress condition were subjected to foot shock stress on days 1 and 3, tail suspension stress on days 2 and 4, and restraint stress on days 3 and 6. All mice were tested on the splash test on day 7, the novelty suppressed feeding test on day 8, the forced swim test on day 9, and the sucrose consumption test on days 8-10 (starting after NSF on day 8). Tissue was collected on day 10.

### Behavioral testing

A battery of behavioral tests was conducted between experimental days 7 and 9 (**Fig. 1**). All mice underwent behavioral testing, and except for the sucrose preference test, all tests were conducted under red light conditions and in designated testing rooms. Animals were acclimatized to the rooms for 1 hour prior to testing.

#### Splash test (ST)

Self-grooming behavior in the ST was used as an index of self care-like and motivational behavior, adapted from previous methods (Hodes et al., 2015; Isingrini et al., 2010). Mice were sprayed three times on the back with a 10% sucrose solution (in tap water), then individually placed in an empty clean cage for 5 mins. The test was filmed, and time spent grooming was scored by an experimenter blinded to condition. Immediately after the self-grooming test, mice were individually housed in clean cages, deprived of food in preparation for the novelty suppressed feeding test, and returned to their colony room. Mice remained individually housed for the remainder of the experiment as per Hodes et al., 2015.

#### Novelty Suppressed Feeding (NSF)

The NSF test was used to assess anxiety-like behavior, and testing was performed according to published protocols (Hodes et al., 2015; Santarelli et al., 2003). Mice were food-deprived overnight, then individually placed in an arena (50×50×20cm) covered with clean bedding and a single pellet of regular chow in the centre. Latency to start feeding was measured as an index of anxiety-like behavior. Mice that did not feed within 10 minutes were removed from the arena and assigned a latency of 600 seconds in the analysis. Bedding was replaced between mice. To account for potential appetite differences between groups, home cage latency to feed and amount of food consumed within 5 minutes were measured.

#### Sucrose Consumption Test (SCT)

The SCT was used to assess anhedonia-like behavior and performed according to previous protocols (Hodes et al., 2015). Mice were first acclimatized to two bottles in the home cage, both filled with tap water. After 24 hours and following the NSF test, the bottles were replaced with two new bottles, one containing tap water and the other 1% sucrose (in tap water). The right-left position was counterbalanced between animals in each condition. The bottles were removed 24 hours later, weighed, then returned to the cage for an additional 24 hours with the right-left position switched for each mouse. Importantly, to avoid hunger-related changes in sucrose consumption, sucrose was presented after food was returned to the home cage following NSF.

#### Forced Swim Test (FST)

The FST was used to assess stress-coping behavior and was conducted according to standard protocols (Can et al., 2012). Mice were individually placed in a 4L glass beaker filled with clean water at 25 ± 0.5°C to a depth of 15cm for a duration of 6 minutes. The test was filmed, and an observer blinded to experimental condition scored two aspects of passive-coping behavior: 1) latency to first bout of immobility and 2) amount of time spent immobile across the duration of the test. Passive-coping behavior was defined as immobility with only movements necessary to remain floating or to keep the head above water.

### Tissue collection and processing

Mice were euthanized via rapid decapitation, and trunk blood was collected into EDTA-coated tubes then centrifuged at 4°C and 1,000 × g. for 15 minutes and plasma was stored at −80°C. Immediately following decapitation, the frontal cortex and hippocampus were micro-dissected on a cold surface, flash-frozen on dry ice, then stored at −80°C. Frontal cortex was collected anterior to the genu of the corpus collosum, and the entire rostral-caudal extent of the hippocampus was collected. Adrenal glands were also collected and weighed immediately.

Electrochemiluminescence immunoassay kits from Meso Scale Discovery (MSD; Rockville, MD) were used according to manufacturer instructions for cytokine, PSD-95, and cell signaling protein measurements. A Sector Imager 2400 (MSD) was used to read the plates, and data was analyzed using the Discovery Workbench 4.0 software from MSD. Frontal cortices and hippocampi from all mice were homogenized individually using an Omni Bead Ruptor (Omni international, Kennesaw, GA) with 200µl and 150µL of cold lysis buffer, respectively. Homogenates were centrifuged at 4°C and 1,000 × g. for 15 minutes, and stored at −80°C. For all assays (cytokine, PSD-95, cell signaling phosphoproteins), frontal cortex and hippocampus values were normalized to total protein concentrations, which were quantified using the Pierce Micro BCA Protein Assay Kit (ThermoFisher Scientific) used according to manufacturer instructions, with samples run in triplicates. Cell signaling proteins and PSD-95 were only quantified in the frontal cortex, as hippocampal tissue remaining after cytokine quantification was insufficient for these assays.

### Cytokine quantification in brain and plasma

V-PLEX proinflammatory Panel 1 kits (Mouse) from MSD (Rockville, MD) were used to measure cytokine concentrations in plasma, frontal cortex, and hippocampus. Plates arrived pre-coated with primary antibodies for the simultaneous quantification of Interferon-γ (IFN-γ), Interleukin-1β (IL-1β), Interleukin-2 (IL-2), Interleukin-4 (IL-4), Interleukin-5 (IL-5), Interleukin-6 (IL-6), chemokine (C-X-C motif) ligand 1 (CXCL1), Interleukin-10 (IL-10), Interleukin-12p70 (IL-12p70), and tumor necrosis factor-α (TNF-α). This panel was chosen because it allows for the quantification of several pro- and anti-inflammatory cytokines as well as the chemokine CXCL1, thus provides a comprehensive view of the cytokine milieu. All samples were run in duplicates, and sample and secondary antibody incubations were performed according to manufacturer instructions. Lower limits of detection (LLOD), which differed between analytes and plates (2 plates total), were as follows: IFN-γ: 0.033-0.076 pg/ml; IL-1β: 0.048-0.069 pg/ml; IL-2: 0.32-0.79pg/ml; IL-4: 0.077-0.091 pg/ml; IL-5: 0.097-0.10 pg/ml; IL-6: 0.83-0.85 pg/ml; CXCL1: 0.15-0.17 pg/ml; IL-10: 0.18-0.62 pg/ml; IL-12p70: 11.8-13.1 pg/ml; TNF-α: 0.082-0.096 pg/ml. In plasma, IL-12p70 was not detectable in 75% of samples, and IL-4 was not detectable in 79% of samples, and thus were not included in analyses.

### Cell signaling phosphoprotein and PSD-95 quantification in the frontal cortex

Cell signaling phosphoproteins (phospho-ERK1/2 (pERK1/2), pMEK1/2, pJNK, pp33, and pSTAT3) and PSD-95 were measured in frontal cortex samples using kits from MSD (Rockville, MD). Primary antibody pre-coated plates were used for sample and secondary antibody incubations according to manufacturer’s instructions, with samples run in duplicates, and results reported after normalization to total protein concentrations in each sample.

### Radioimmunoassays for hormone quantification in plasma

Plasma corticosterone concentrations from samples collected at euthanasia were quantified using a commercially available radioimmunoassay kit used in accordance with manufacturers’ instructions (corticosterone double-antibody radioimmunoassay kit, MP biomedicals, Solon, OH). To avoid handling-induced elevations in circulating glucocorticoids, trunk blood samples were collected within 2 minutes of the experimenter entering the colony room. The inter- and intra-assay coefficients of variation were below 10%.

An Ultra-Sensitive Estradiol Radioimmunoassay kit (Beckman Coulter, Prague, Czech Republic) was used according to manufacturers’ instructions to measure 17β-estradiol concentrations in plasma samples collected at euthanasia. Assay sensitivity is 2.2pg/ml, and average inter- and intra-assay coefficients of variation were below 10%.

### Vaginal cytology for estrous cycle assessment

Estrous cycle stage can affect behavior, measures of neural plasticity, and immune function (Beagley and Gockel, 2003; Meziane et al., 2007; Woolley et al., 1990), thus vaginal lavage samples were obtained on behavioral testing days, and estrous stage was determined according to previous methods (Cora et al., 2015). Plasma was collected on the last day for estradiol quantification. We expected irregular estrous cycling in at least a proportion of sham-operated mice as they were middle-aged at testing. Mice in persistent diestrus, persistent estrus, or that displayed abnormalities in the length or order of estrous cycle stages were all classified as irregularly cycling, as we have done previously (Galea et al., 2018). Ovariectomized mice were also lavaged to control for potential effects of the procedure.

### Statistical analyses

All statistical analyses were performed using Statistica software (Tulsa, OK). Behavioral measures, cytokine concentrations, IL-6:IL-10 ratio, corticosterone concentrations, PSD-95, and cell signaling protein levels were each analyzed using factorial analysis of variance (ANOVA), with ovarian status (OVX, Sham) and stress condition (stress, non-stress) as the between-subject factors. Body mass was used as a covariate when analyzing sucrose consumption, immobility in the forced swim test, self-grooming behavior in the splash test, and adrenal mass. Estrous cycle stage (proestrus, non-proestrus) was used as a covariate in behavioral analyses. 17β-estradiol concentrations were used as a covariate in corticosterone, cytokine, cell signaling, and PSD-965 analyses. Covariate effects are only mentioned when significant. Post-hoc analyses utilized Newman-Keul’s comparisons and any *a priori* comparisons were subjected to a Bonferroni correction. A Chi-square test was used to compare the frequency of mice that had regular estrous cycles between sham-operated groups. To assess whether cycle regularity affected outcomes in sham-operated mice, we analyzed data from sham-operated mice separately using ANOVAs with cycle regularity (regular, irregular) and stress condition (stress, non-stress) as the between-subject factors. Specifically, we analyzed all behavioral outcomes and key cytokines and cell signaling phosphoproteins that were found to be significantly affected in initial ANOVAs. However, we did not find any significant effects of cycle regularity nor of the interaction between stress condition and cycle regularity (all p’s >0.2), therefore these analyses are not further reported in the results section. Pearson’s correlations were performed between variables of interest. Outliers that fell more than 2.5 standard deviations away from the mean were removed from analyses. Principal Component Analyses (PCA) were used to reduce the cytokine data into a smaller number of uncorrelated variables and to obtain information about the amount of variance accounted for by potential cytokine networks within the data. PCAs were followed by ANOVAs on individual principal component scores as we have done previously (Eid et al., 2019a), to assess the effects of ovarian status and stress exposure on cytokine networks.

## Results

### Body mass was significantly reduced across days of stress exposure regardless of ovarian status

Long-term ovariectomy significantly increased body mass (F(1, 24)=22.72, p<0.0001), therefore to investigate the effects of sub-chronic stress exposure we calculated body mass as a percentage of mass on experimental day 1. Body mass percentage decreased significantly across the 6 days of stress exposure (F(4, 96)=32.967, p<0.0001; stress condition by day interaction; **Fig. 2**). Specifically, in stress-exposed groups, there was a significant decline on day 3 relative to day 2, and on every day thereafter (p’s <0.006). Body mass percentage was also significantly different between stress and non-stress groups on days 3-6 (p’s <0.007). There were also significant main effects of stress condition and day (p’s<0.0001), a trend toward a significant day by ovarian status interaction (p=0.064), but no other significant main effects or interactions (p’s >0.095).

**Figure 2.**
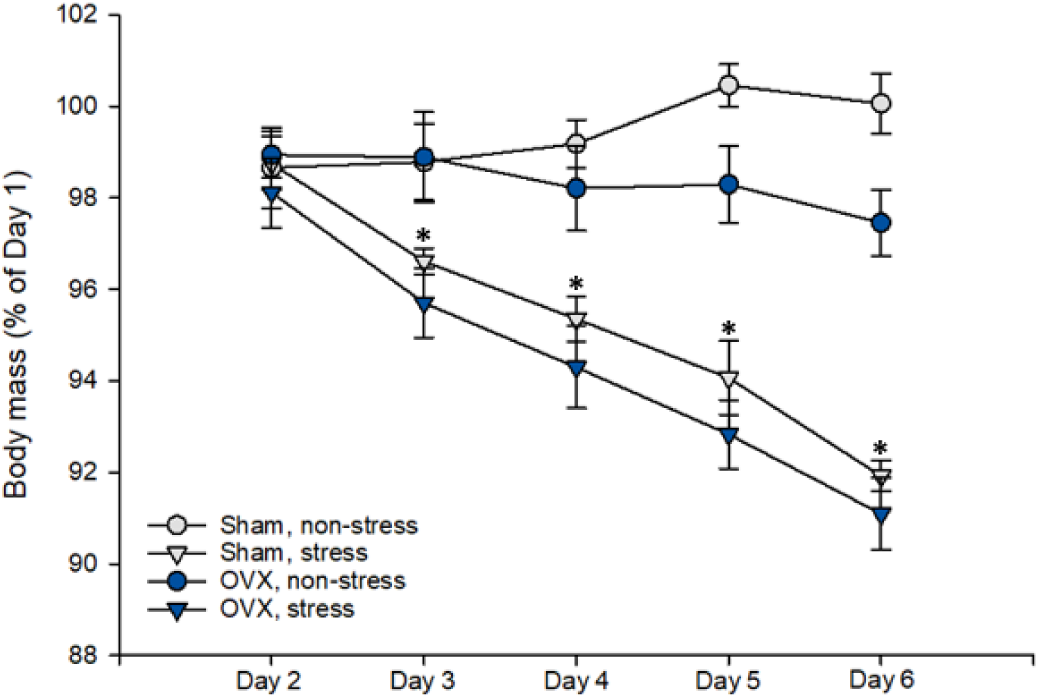
Body mass across days, shown as a percentage of body mass on experimental day 1. Regardless of ovarian status, body mass decreased significantly across days with sub-chronic stress exposure; * indicates p<0.007, significantly different from day-matched non-stress groups, and stress groups on prior day/s of stress exposure. n = 7/group. Data in means ± standard error of the mean. OVX, ovariectomized.

### Long-term ovariectomy reduced circulating 17β-estradiol concentrations but stress exposure did not affect 17β-estradiol concentrations or estrous cycling

Plasma 17β-estradiol concentrations were significantly reduced by long-term ovariectomy (F(1, 18)=9.20, p=0.007; main effect of ovarian status). Stress condition did not affect 17β-estradiol concentrations, nor did the interaction of stress condition and ovarian status (all p’s >0.2; **Table 1**). These results do not change when analyzing estrous cycle regularity as a covariate: a significant main effect of ovarian status remains (p = 0.018), but there was no significant main effect of the covariate, and no significant effect of stress condition nor a stress condition by ovarian status interactions (p’s >0.2). Within sham-operated groups, there was no significant difference in the frequency of mice that had regular estrous cycles (χ^2^(1) = 0.31, p = 0.58; **Table 1**). Further, there were no significant differences in 17β-estradiol concentrations between regularly vs. irregularly cycling mice, regardless of stress condition (all p’s >0.4). It should be noted that 17β-estradiol concentrations in sham-operated groups were considerably lower than what would be expected in young-adult intact mice (∼20-60pg/ml across estrous cycle (Walmer et al., 1992)), indicating our that sham-operated groups were in a transitional state to reproductive senescence.

**Table 1.** Plasma 17β-estradiol concentrations from trunk blood at decapitation and percentage of sham-operated mice with regular estrous cycles.

| Group | $17\beta$-estradiol (pg/ml) | % of mice regularly cycling |
| --- | --- | --- |
| Sham, non-stress | $10.34 \pm 1.85$ | 42.86 |
| Sham, stress | $10.32 \pm 2.20$ | 28.57 |
| OVX, non-stress | $4.30 \pm 0.96$ | N/A |
| OVX, stress | $7.52 \pm 0.71$ | N/A |

Data in means ± standard error of the mean. $n = 6$ - $7$ /group. OVX, ovariectomized.

### Basal plasma corticosterone concentrations and adrenal mass did not differ significantly between groups

Basal plasma corticosterone concentrations did not differ significantly between groups (all p’s >0.3; **Table 2**), but there was a significant covariate effect of plasma 17β-estradiol concentrations (p = 0.027). Similarly, adrenal mass was not significantly affected by ovarian status, stress exposure, nor their interaction (p’s>0.3; **Table 2**).

**Table 2.** Basal plasma corticosterone concentrations and adrenal mass.

| Group | CORT (pg/ml) | Adrenal mass (mg) |
| --- | --- | --- |
| Sham, non-stress | $42.48 \pm 8.99$ | $6.99 \pm 0.66$ |
| Sham, stress | $33.79 \pm 6.70$ | $8.06 \pm 0.53$ |
| OVX, non-stress | $31.92 \pm 6.88$ | $8.50 \pm 0.88$ |
| OVX, stress | $39.58 \pm 8.62$ | $8.16 \pm 0.69$ |

Data in means ± standard error of the mean. $n = 6$ - $7$ /group. OVX, ovariectomized; CORT, corticosterone.

### Ovariectomy reduced latency to immobility under non-stress conditions, and sub-chronic stress exposure reduced latency to immobility in sham-operated mice only

Under non-stress conditions, ovariectomy significantly decreased latency to immobility in the FST (p=0.01; *a priori* comparisons; ovarian status by stress condition interaction: F(1, 18)=1.8372, p=0.19). Further, exposure to sub-chronic stress significantly decreased the latency to immobility in sham-operated (p=0.012; **Fig. 3A**) but not in ovariectomized mice (p=0.54). There was also a significant main effect of stress condition (p=0.025) but not of ovarian status (p=0.22). The percentage of time spent immobile in FST did not significantly differ between groups (all p’s >0.09; **Table 3**)

**Figure 3.**
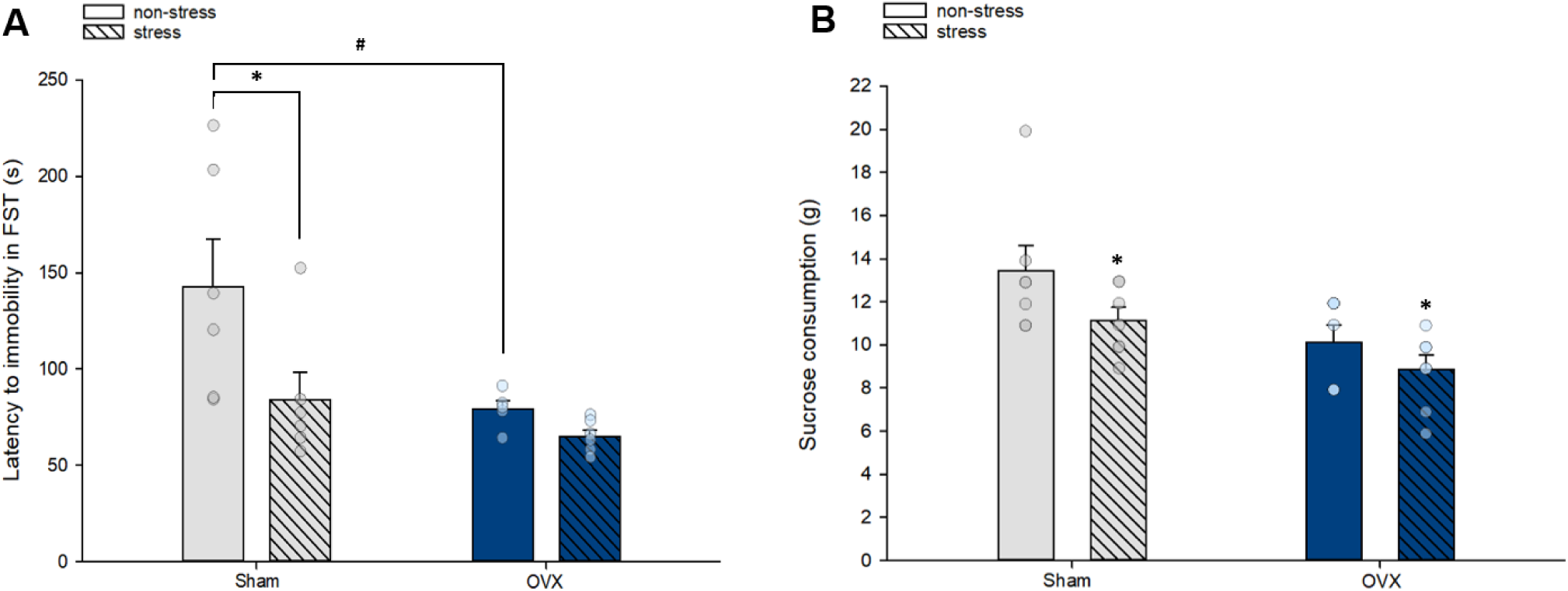
Latency to first bout of immobility in the Forced Swim Test and sucrose consumption in the Sucrose Consumption Test. **(A)** Ovariectomy significantly reduced latency to immobility under non-stress conditions, and sub-chronic stress exposure reduced latency to immobility in sham-operated mice only; **#** indicates p=0.01 and *indicates p=0.012. **(B)** Stress exposure decreased sucrose consumption, regardless of ovarian status, and there was a trend for ovariectomy to reduce sucrose consumption; * indicates p=0.038, main effect of stress condition. FST = forced swim test; OVX = ovariectomy. n = 6-7/group. Data in means + standard error of the mean.

**Table 3.**
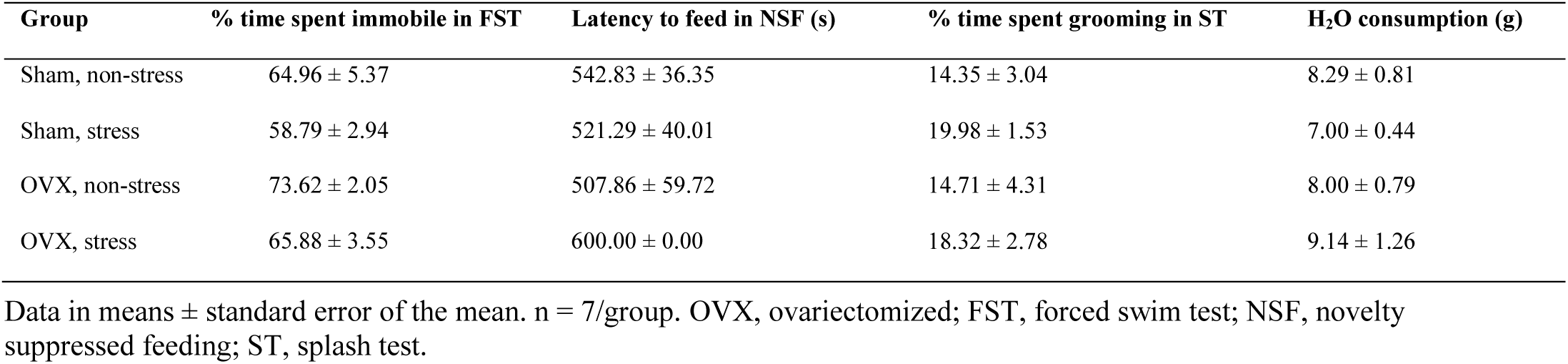
Behavioral data.

| Group | % time spent immobile in FST | Latency to feed in NSF (s) | % time spent grooming in ST | H<sub>2</sub>O consumption (g) |
| --- | --- | --- | --- | --- |
| Sham, non-stress | 64.96 ± 5.37 | 542.83 ± 36.35 | 14.35 ± 3.04 | 8.29 ± 0.81 |
| Sham, stress | 58.79 ± 2.94 | 521.29 ± 40.01 | 19.98 ± 1.53 | 7.00 ± 0.44 |
| OVX, non-stress | 73.62 ± 2.05 | 507.86 ± 59.72 | 14.71 ± 4.31 | 8.00 ± 0.79 |
| OVX, stress | 65.88 ± 3.55 | 600.00 ± 0.00 | 18.32 ± 2.78 | 9.14 ± 1.26 |

Data in means ± standard error of the mean. $n = 7$ /group. OVX, ovariectomized; FST, forced swim test; NSF, novelty suppressed feeding; ST, splash test.

### Sub-chronic stress exposure decreased sucrose consumption regardless of ovarian status

Stress exposure significantly reduced sucrose consumption (F(1, 23)=4.87, p=0.038; main effect of stress condition; **Fig. 3B**). There was also a trend for ovariectomy to reduce sucrose consumption (F(1, 23)=3.98, p=0.067), but no other significant effects (p’s>0.4). In comparison, water consumption was not significantly affected by body mass, ovarian status, stress condition, nor their interaction (all p’s >0.4; **Table 3**).

### Sub-chronic stress exposure and ovarian status did not significantly affect behavior in the splash and novelty suppressed feeding tests

Anxiety-like behavior as measured in the novelty suppressed feeding test and self-grooming behavior in the splash test were not significantly altered by ovariectomy, sub-chronic stress exposure, nor their interaction (all p’s >0.1; **Table 3**).

### Principal component (PC) analyses in cytokine data

#### Frontal cortex: PC1 scores were reduced by long-term ovariectomy, and PC2 scores were increased by stress exposure in sham-operated mice only

The model generated 3 principal components, accounting for 81.8% of the variance within the frontal cortex cytokine dataset. Variance explained by principal component 1 (PC1) = 52.8%, PC2 = 17.1 %, and PC3 = 11.9%. Factor loadings are shown in **Table 4**. ANOVA results reveal that ovariectomy significantly reduced PC1 scores (F(1, 24)=5.23, p=0.031; main effect of ovarian status; **Fig. 4A**), but there was no significant main effect of stress condition, nor an ovarian status by stress condition interaction (p’s >0.6). As such we will refer to PC1 as “Ovarian hormone sensitive” cytokines. Stress significantly increased PC2 scores in sham (p = 0.008), but not ovariectomized mice (p = 0.51; *a priori* comparisons, ovarian status by stress condition interaction: F(1, 24)=2.526, p<0.13; **Fig. 4B**). There was also a significant main effect of stress condition (p = 0.019), but not of ovarian status (p = 0.55). For PC3 scores, there were no significant main effects of ovarian status or stress condition, nor a significant interaction (p’s >0.19). Thus, in the frontal cortex we identify 8 cytokines that were proportionally more sensitive to ovarian hormone status based on component loadings(IL-2, IL-4 IL5, IL-6, IL-10, TNF-α, IL-12p70 and the chemokine CXCL1) and 3 cytokines sensitive to both stress and ovarian hormone status (IL-1β, TNF-α and IL-4).

**Figure 4.**
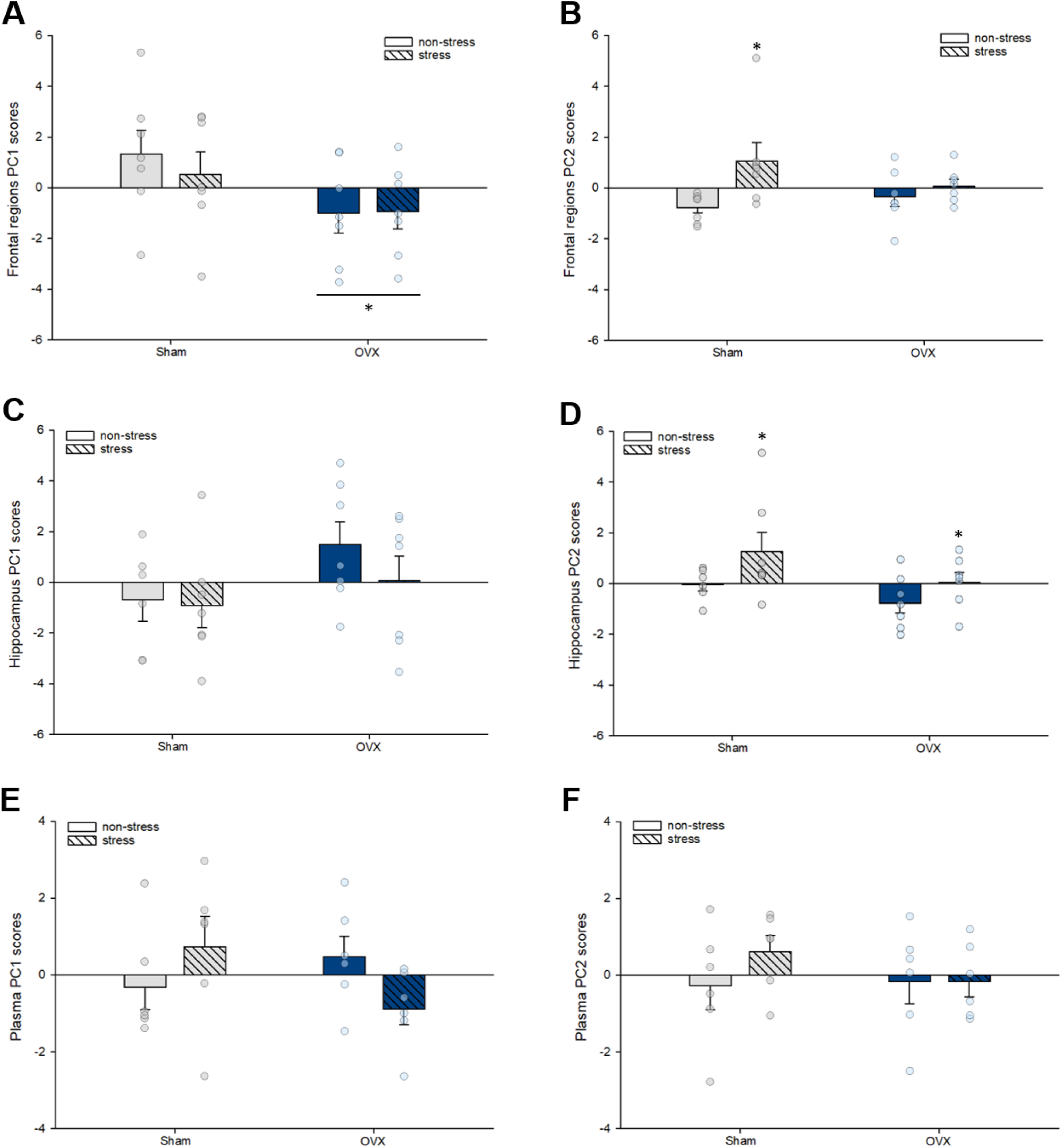
Principal component analyses scores in cytokine data. **(A)** Long-term ovariectomy significantly decreased Principal Component (PC) 1 scores in frontal cortex; * indicates p=0.031, main effect of ovarian status. **(B)** Exposure to sub-chronic stress increased PC2 scores in the frontal cortex in sham-operated mice only; * indicates p=0.008, significantly higher than non-stress sham-operated group. **(C)** There was a weak trend (p=0.096) for ovariectomy to increase PC1 scores in the hippocampus. **(D)** Sub-chronic stress exposure increased PC2 scores in the hippocampus, regardless of ovarian status, and there was a trend for ovariectomy (p=0.064) to reduce PC2 scores; * indicates p=0.047, main effects of stress condition. There were no significant group differences in PC1 or PC2 scores in plasma **(E-F)**. PC = principal component; OVX = ovariectomy. n = 6-7/group. Data in means + standard error of the mean.

**Table 4.** Principal component analyses loading table

|  | Frontal cortex |  |  | Hippocampus |  |  | Plasma |  |
| --- | --- | --- | --- | --- | --- | --- | --- | --- |
| Cytokine | PC1 | PC2 | PC3 | PC1 | PC2 | PC3 | PC1 | PC2 |
| IFN-γ | 0.22 | -0.30 | **0.91** | **0.85** | -0.03 | -0.24 | **0.59** | 0.17 |
| IL-10 | **0.93** | -0.21 | 0.25 | **0.90** | -0.19 | 0.09 | -0.25 | 0.25 |
| IL-1β | 0.22 | **0.85** | 0.36 | 0.45 | **0.58** | -0.35 | **0.59** | **-0.67** |
| IL-2 | **0.90** | 0.00 | -0.08 | **0.89** | -0.19 | -0.17 | **0.73** | -0.25 |
| IL-5 | **0.87** | -0.23 | 0.08 | **0.65** | -0.30 | 0.71 | 0.24 | **0.65** |
| IL-6 | **0.77** | 0.10 | -0.09 | **0.88** | -0.007 | 0.19 | **0.67** | 0.24 |
| CXCL1 | **0.73** | 0.35 | -0.30 | 0.17 | **1.13** | **0.44** | **0.65** | 0.03 |
| TNF- $\alpha$ | **0.75** | **0.47** | 0.18 | **0.64** | **0.54** | -0.31 | 0.26 | **0.67** |
| IL-12p70 | **0.90** | -0.11 | -0.17 | **0.92** | -0.16 | 0.064 | N/A | N/A |
| IL-4 | **0.49** | **-0.66** | 0.04 | **0.89** | -0.11 | 0.09 | N/A | N/A |

High loadings indicated in bold.

#### Hippocampus: Stress exposure increased PC2 scores regardless of ovarian status, and there were trends for long-term ovariectomy to increase PC1 and decrease PC2 scores

The model generated 3 principal components, accounting for 81.4% of the variance within the hippocampus cytokine dataset. Variance explained by principal component 1 (PC1) = 58.0%, PC2 = 14.8%, and PC3 = 8.6%. Factor loadings are shown in **Table 4**. ANOVAs reveal a weak trend for ovariectomy to increase PC1 scores (F(1, 23)=3.02, p=0.096; main effect of ovarian status; **Fig. 4C**), and no significant main effect of stress condition, nor an ovarian status by stress condition interaction (p’s >0.3). Stress significantly increased PC2 scores regardless of ovarian status (F(1, 23)=4.40, p=0.047; **Fig. 4D**), and there was a trend toward significance for ovariectomy to decrease PC2 scores (F(1, 23)=3.67, p=0.068), but no significant stress condition by ovarian status interaction (p > 0.6). For PC3 scores, there were no significant main effects of ovarian status or stress condition, nor a significant interaction (p’s >0.3). Thus, these analyses in the hippocampus identify 8 cytokines that were sensitive to ovarian hormone status based on component loadings (IL-2, IL-4 IL5, IL-6, IL-10, TNF-α, IL-12p70, IFN-γ) and 3 cytokines sensitive to stress (IL-1β, TNF-α and the chemokine CXCL1).

#### Plasma: PC scores did not differ significantly between groups

The model generated 2 principal components, accounting for 48.2% of the variance within the plasma cytokine dataset. Variance explained by principal component 1 (PC1) = 29.0%, and PC2 = 19.2%. Factor loadings are shown in **Table 4**. ANOVAs reveal no significant main effects of stress condition or ovarian status, nor a significant interaction for PC1 or PC2 scores (p’s > 0.06; **Fig. 4E-F**).

### Individual cytokine ANOVAs

#### Long-term ovariectomy reduced the concentrations of several cytokines in the frontal cortex

Long-term ovariectomy significantly reduced the concentrations of the following cytokines/chemokine in the frontal cortex: TNF-α (F(1, 24)=5.41, p=0.029; **Fig. 5A**); CXCL1 (F(1, 24)=4.44, p=0.046; **Fig. 5B**); IL-10 (F(1, 22)=9.80, p=0.0049; **Fig. 5C**); IL-12p70 (F(1, 24)=6.40, p=0.018; **Fig. 5D**), IFN-γ (F(1, 23)=5.89, p=0.023; **Fig. 5E**), and IL-6 (F(1, 24)=4.16, p=0.05; **Fig. 5F**). For each of these cytokines, there was no significant main effect of stress condition, and no stress condition by ovarian status interaction (p’s > 0.2).

**Figure 5.**
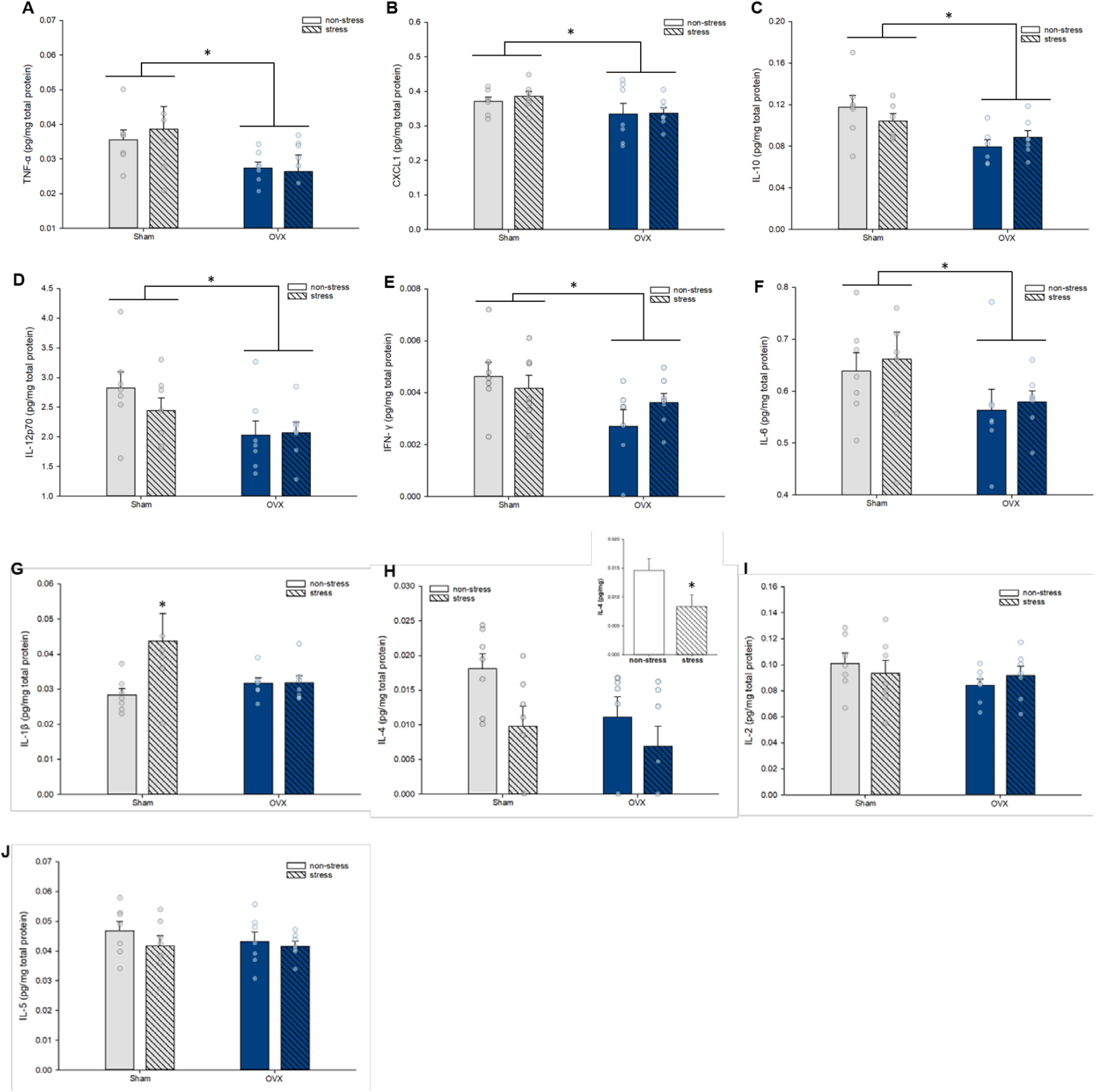
Cytokine concentrations in the frontal cortex, normalized by total protein concentrations. Long-term ovariectomy reduced the concentrations of TNF-α **(A)** CXCL1 **(B)**, IL-10 **(C)**, IL-12p70 **(D)**, IFN-γ **(E)**, and IL-6 **(F)** regardless of stress exposure; * indicates significant main effects of ovarian status (p’s <0.05). **(G)** Stress exposure increased IL-1β in the frontal cortex of sham-operated mice only; * indicates p = 0.018, significantly higher than non-stress sham group. **(H)** Stress exposure significantly reduced IL-4; inset graph depicts main effect of stress condition, * indicates p = 0.031, significantly lower than non-stress. Concentrations of IL-2 **(I)** and IL-5 **(J)** did not differ significantly between groups. OVX, ovariectomized; IL, interleukin; CXCL1, chemokine (C-X-C motif) ligand 1; TNF-α, tumor necrosis factor-α; IFN-γ, Interferon-γ. n = 6-7/group. Data in means + standard error of the mean.

#### Sub-chronic stress exposure significantly increased IL-1β in the frontal cortex of sham-operated mice only

Exposure to stress increased IL-1β concentrations in the frontal cortex of sham-operated (p=0.018, a priori comparison) but not ovariectomized mice (p=0.98; F(1, 24)=3.1706, p=0.088; stress condition by ovarian status interaction; **Fig. 5G**). There was also a trend toward a significant main effect of stress exposure to increase IL-1β (p= 0.082), but no significant main effect of ovarian status (p = 0.32).

#### Sub-chronic stress exposure significantly decreased IL-4 in the frontal cortex

Stress exposure significantly reduced IL-4 concentrations in the frontal cortex (F(1, 24)=5.28, p=0.031; main effect of condition; **Fig. 5H**). This effect was driven by a larger decrease in sham-operated mice, as *a priori* comparisons reveal a trend toward a significant difference between sham-operated groups (p = 0.041) but not between ovariectomized groups (p = 0.28). There was also a trend toward significance for ovariectomy to reduce IL-4 (p = 0.085), but no significant stress condition by ovarian status interaction (p = 0.46). There were no significant effects of stress condition or ovarian status, nor an interaction to affect IL-2 (P’s > 0.25; **Fig. 5I**) or IL-5 (p’s>0.25; **Fig. 5J**) concentrations in the frontal cortex.

#### Long-term ovariectomy increased IL-2, IL-5, and IL-6 concentrations in the hippocampus

Long-term ovariectomy significantly increased IL-2 (F(1, 22)=8.92, p<0.007; **Fig. 6A**), IL-5 (F(1, 22)=8.37, p=0.008; **Fig. 6B**), and IL-6 (F(1, 23)=5.80, p=0.024; **Fig. 6C**) concentrations in the hippocampus. For each of these cytokines, there was no significant main effect of stress condition, nor a stress condition by ovarian status interaction (all p’s >0.18).

**Figure 6.**
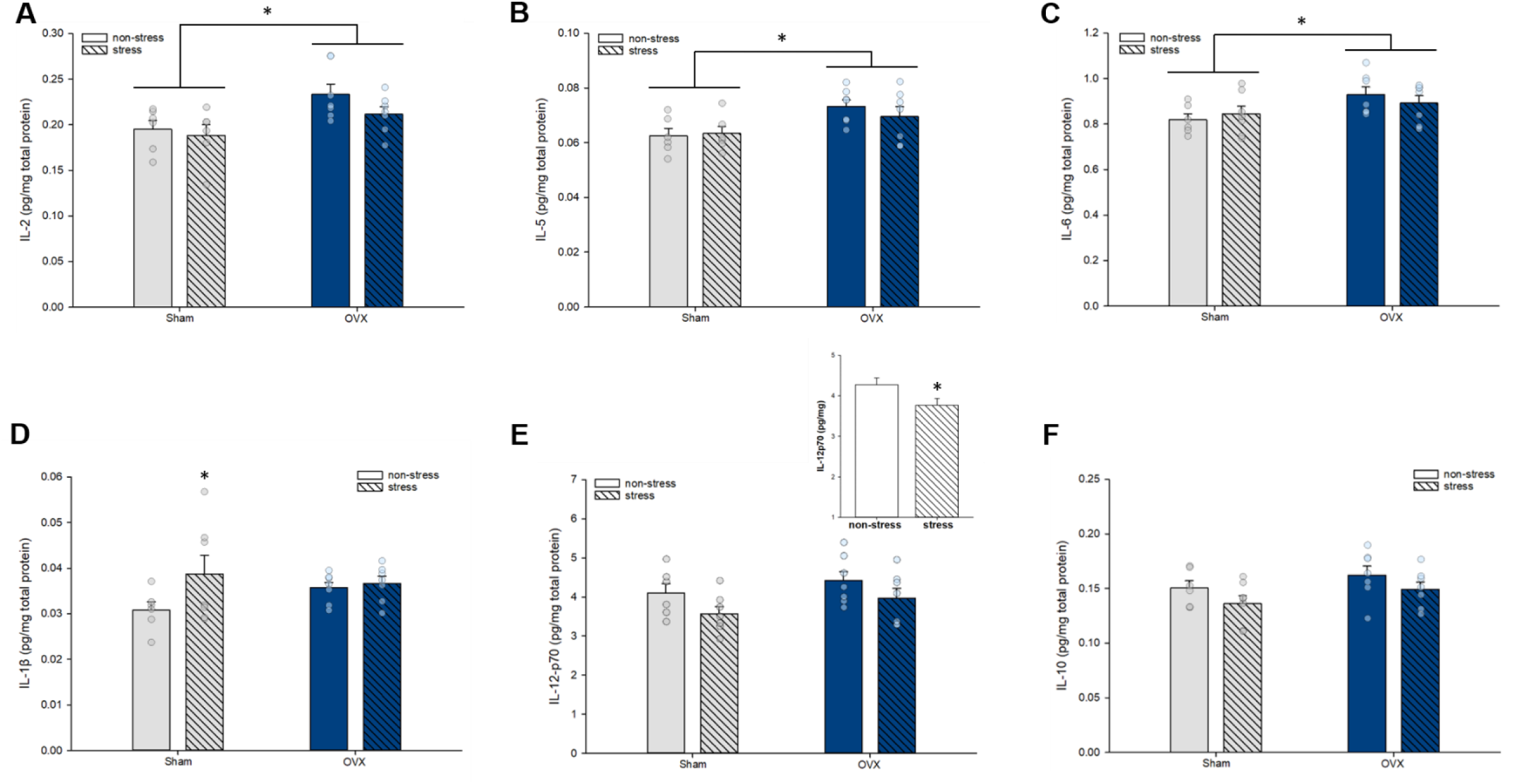
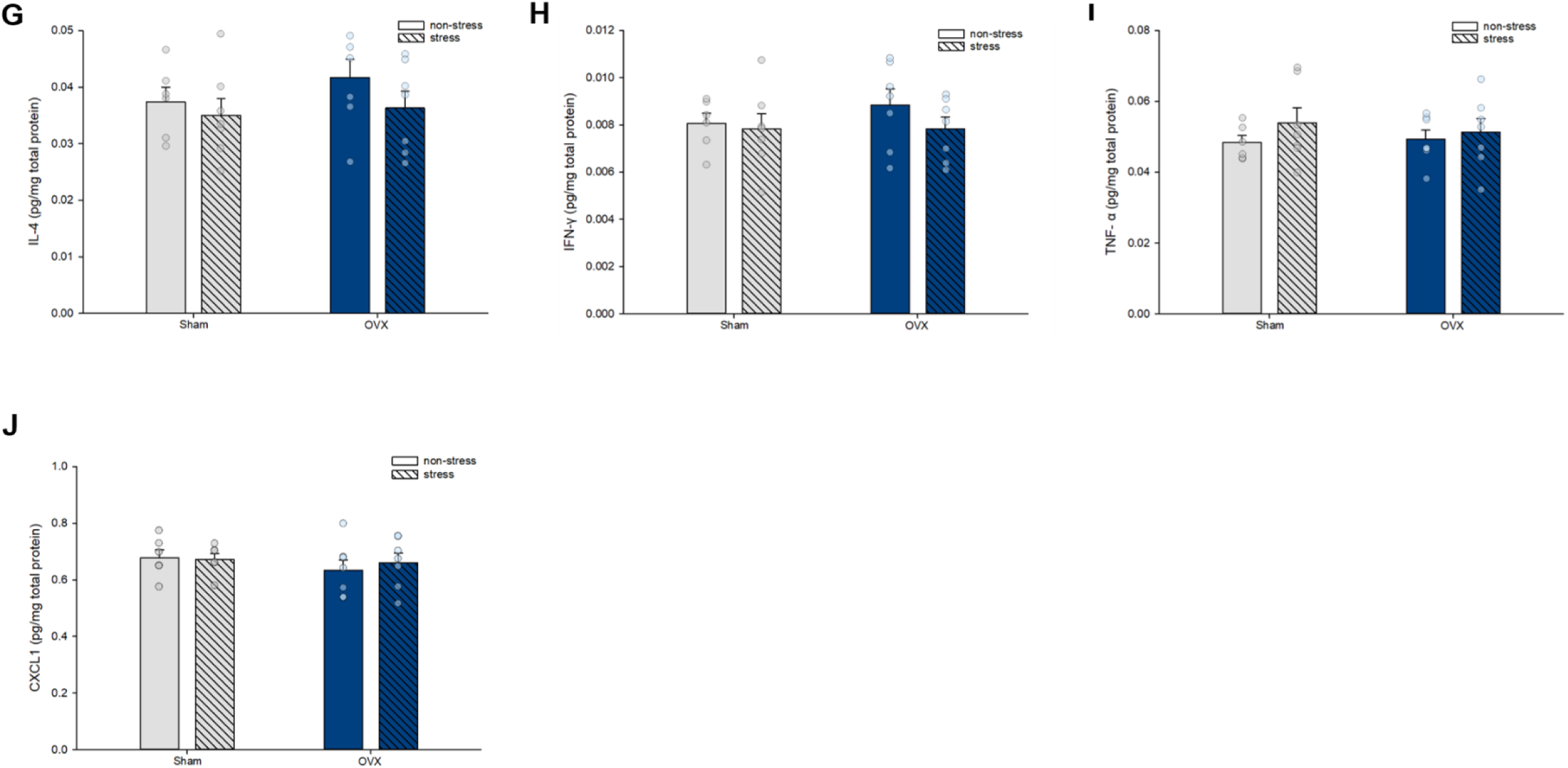
Cytokine concentrations in the hippocampus, normalized by total protein concentrations. Long-term ovariectomy significantly increased concentrations of IL-2 **(A)**, IL-5 **(B)**, and IL-6 **(C)** in the hippocampus; * indicates p<0.03, main effect of ovarian status**. (D)** Stress exposure increased IL-1β in the hippocampus of sham-operated mice only; * indicates p = 0.04, significantly higher than non-stress sham group. **(E)** Sub-chronic stress exposure significantly decreased IL-12p70 concentrations, regardless of ovarian status; inset graph depicts main effect of stress condition, * indicates p<0.05, significantly lower than non-stress. Hippocampal concentrations of IL-10 **(F)**, IL-4 **(G)**, IFN-γ **(H)**, TNF-α **(I)**, and CXCL1 **(J)** did not significantly differ between groups. OVX, ovariectomized; IL, interleukin; CXCL1, chemokine (C-X-C motif) ligand 1; TNF-α, tumor necrosis factor-α; IFN-γ, Interferon-γ. n = 6-7/group. Data in means + standard error of the mean.

#### Sub-chronic stress exposure increased IL-1β in the hippocampus of sham-operated mice only, and decreased IL-12p70 concentrations regardless of ovarian status

Sub-chronic stress exposure increased IL-1β concentrations in the hippocampus of sham-operated (p=0.04) but not ovariectomized mice (p=0.78; F(1, 23)=1.82, p=0.19; *a priori* comparison; stress condition by ovarian status interaction; **Fig. 6D**). There were no significant main effects of stress condition (p=0.09) or ovarian status (p>0.5). Regardless of ovarian status, exposure to sub-chronic stress significantly decreased hippocampal IL-12p70 concentrations (F(1, 23)=4.38, p=0.048; **Fig. 6E**). There was no significant main effect of ovarian status nor an ovarian status by stress condition interaction (p’s >0.1). There was also a weak trend for stress exposure to reduce IL-10 (p=0.082; **Fig. 6F**), but no significant group differences for hippocampal IL-4 (p’s > 0.2; **Fig. 6G**), IFN-γ (p’s > 0.29; **Fig. 6H**), TNF-α (p’s > 0.27; **Fig. 6I**), or CXCL1 (p’s > 0.3; **Fig. 6J**).

#### Ovariectomy significantly increased plasma CXCL1 concentrations

Ovariectomy significantly increased CXCL1 concentrations in plasma (F(1,20)=6.62, p=0.018; main effect of ovarian status; **Fig 7A**), but there was no significant main effect of stress condition (p=0.23) nor an ovarian status by stress condition interaction (p = 0.53). There were trends toward significance for an ovarian status by stress condition interaction to affect plasma IL-10 (F(1, 20)=3.85, p=0.064; **Fig. 7B**) and IL-6 (F(1, 19)=3.2520, p=0.087; **Fig. 7C**) concentrations, but no significant main effects of ovarian status nor stress condition in both cases (p’s > 0.19). There were no significant main effects of stress condition or ovarian status nor a significant interaction for plasma IL-5 (all p’s >0.29; **Fig. 7D**), TNF-α (p’s > 0.14; **Fig. 7E**), IFN-γ (p’s > 0.4; **Fig. 7F**), IL-2 (p’s > 0.6; **Fig. 7G**), or IL-1β (p’s > 0.2; **Fig. 7H**).

**Figure 7.**
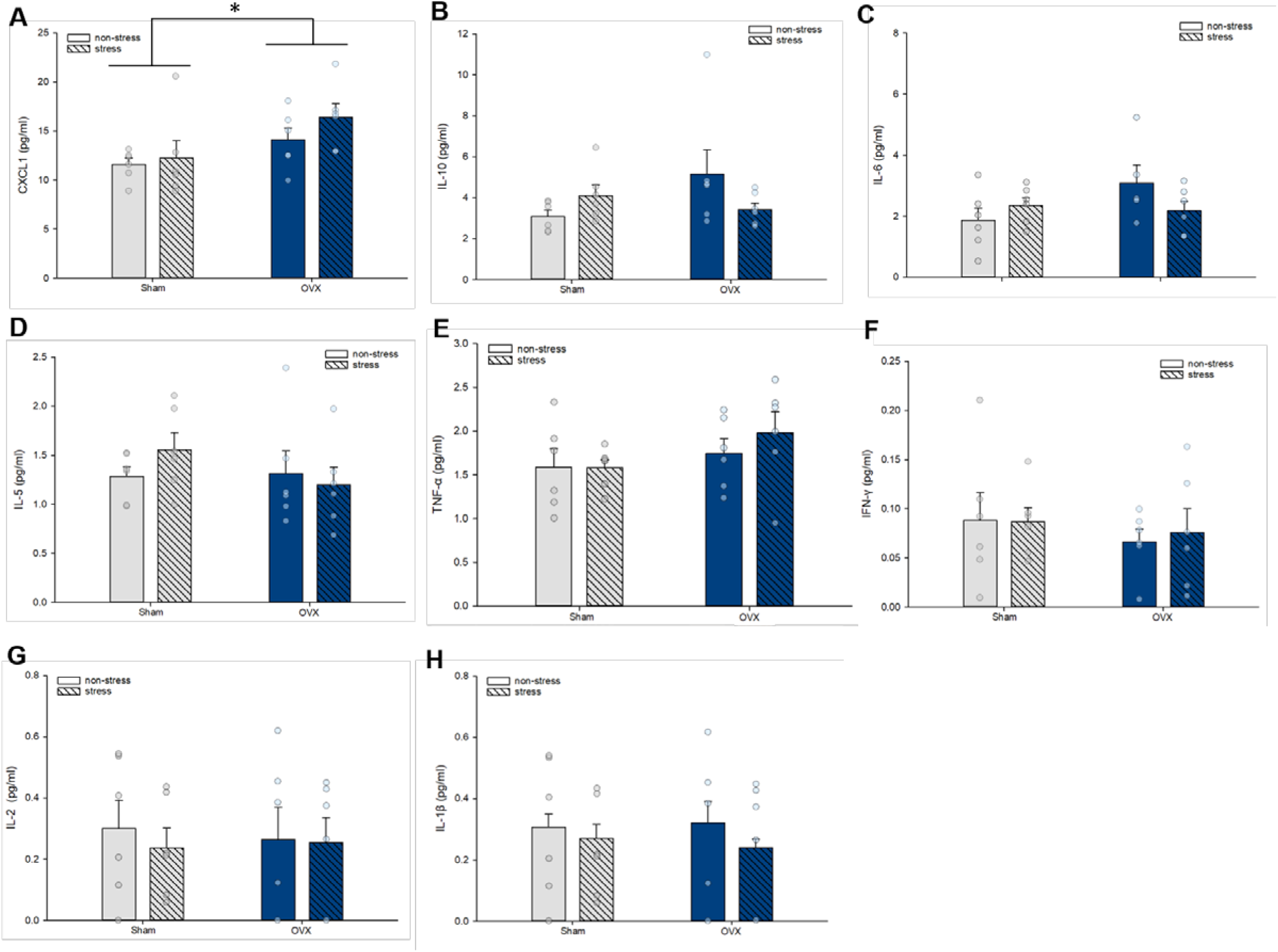
Cytokine concentration in plasma. **(A)** Ovariectomy significantly increased CXCL1 concentrations in plasma; * indicates p= 0.018, main effect of ovarian status. There were no significant group differences in plasma concentrations of IL-10 **(B)**, IL-6 **(C)**, IL-5 **(D)**, TNF-α **(E)**, IFN-γ **(F)**, IL-2 **(G)**, or IL-1β **(H)**. OVX, ovariectomized; IL, interleukin; CXCL1, chemokine (C-X-C motif) ligand 1; TNF-α, tumor necrosis factor-α; IFN-γ, Interferon-γ. n = 6/group. Data in means + standard error of the mean.

#### Sub-chronic stress exposure increased IL-6:IL-10 ratio in the frontal cortex and hippocampus in sham-operated mice only

IL-6:IL-10 ratios were analyzed as an indicator of pro- to anti-inflammatory cytokine balance as previously described (de Brito et al., 2016; Wood et al., 2015). In both brain regions, sub-chronic stress exposure significantly increased IL-6:IL-10 ratio in sham-operated mice only (frontal cortex: p = 0.033; ovarian status by stress condition interaction: F(1, 22)=4.72, p=0.041, **Fig 8A**; hippocampus: p = 0.015; *a priori* comparisons; stress condition by ovarian status interaction: F(1, 23)=1.9553, p=0.17, **Fig. 8B**). In the hippocampus, there was also a significant main effect of condition, with stress increasing the ratio (p=0.025), but not of ovarian status (p=0.87), and there were no significant main effects in the frontal cortex (p’s >0.12). Finally, under non-stress conditions, ovariectomy significantly increased IL-6:IL-10 ratio in the frontal cortex only (p = 0.014; **Fig. 8A**). For plasma IL-6:IL-10 ratios, there were no significant main effects, nor a significant interaction (all p’s >0.3).

**Figure 8.**
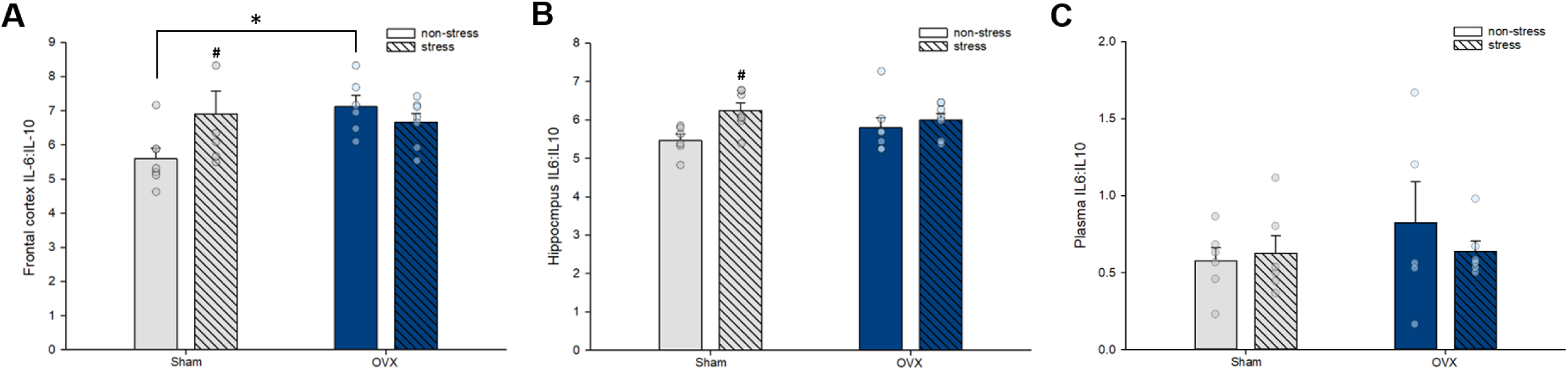
Interleukin-6 to interleukin-10 ratio in the frontal cortex **(A)**, hippocampus **(B)**, and plasma **(C)**. **(A)** Ovariectomy significantly increased IL-6:IL-10 ratio in the frontal cortex, and sub-chronic stress exposure increased IL-6:IL-10 ratio in sham-operated mice only. * indicates p= 0.014 and # indicates p=0.033, significantly higher than non-stress sham group. **(B)** sub-chronic stress exposure increased IL-6:IL-10 ratio in the hippocampus in sham-operated mice only. # indicates p=0.015, significantly higher than non-stress sham group. **(C)** There were no significant group differences in plasma IL-6:IL-10 ratio. OVX, ovariectomized; IL, interleukin. n = 6-7/group. Data in means + standard error of the mean.

#### pERK1/2 and pMEK1/2 were significantly reduced by long-term ovariectomy and the effects of stress were dependent on ovarian status

Ovariectomy significantly decreased pERK1/2 and pMEK1/2 expression in the frontal cortex under non-stress conditions, relative to sham-operated mice (both p’s = 0.032). Sub-chronic stress exposure significantly decreased pERK1/2 expression in sham-operated mice (p= 0.023), but not in ovariectomized mice (p=0.35; ovarian status by stress condition interaction: F(1, 24)=5.68, p=0.025, **Fig. 9A**). Conversely, sub-chronic stress exposure significantly increased pMEK1/2 expression in ovariectomized mice (p= 0.042), but not in sham-operated mice (p = 0.31; significant ovarian status by stress condition interaction F(1, 24)=5.065, p=0.034, **Fig 9B**). A similar, albeit non-significant trend was observed for pp38 expression (ovarian status by stress condition interaction (F(1, 24)=3.844, p=0.062; **Fig. 9C**), and there were no significant main or interaction effects for pJNK or pSTAT3 expression (all p’s >0.14; **Fig 9D-E**).

**Figure 9.**
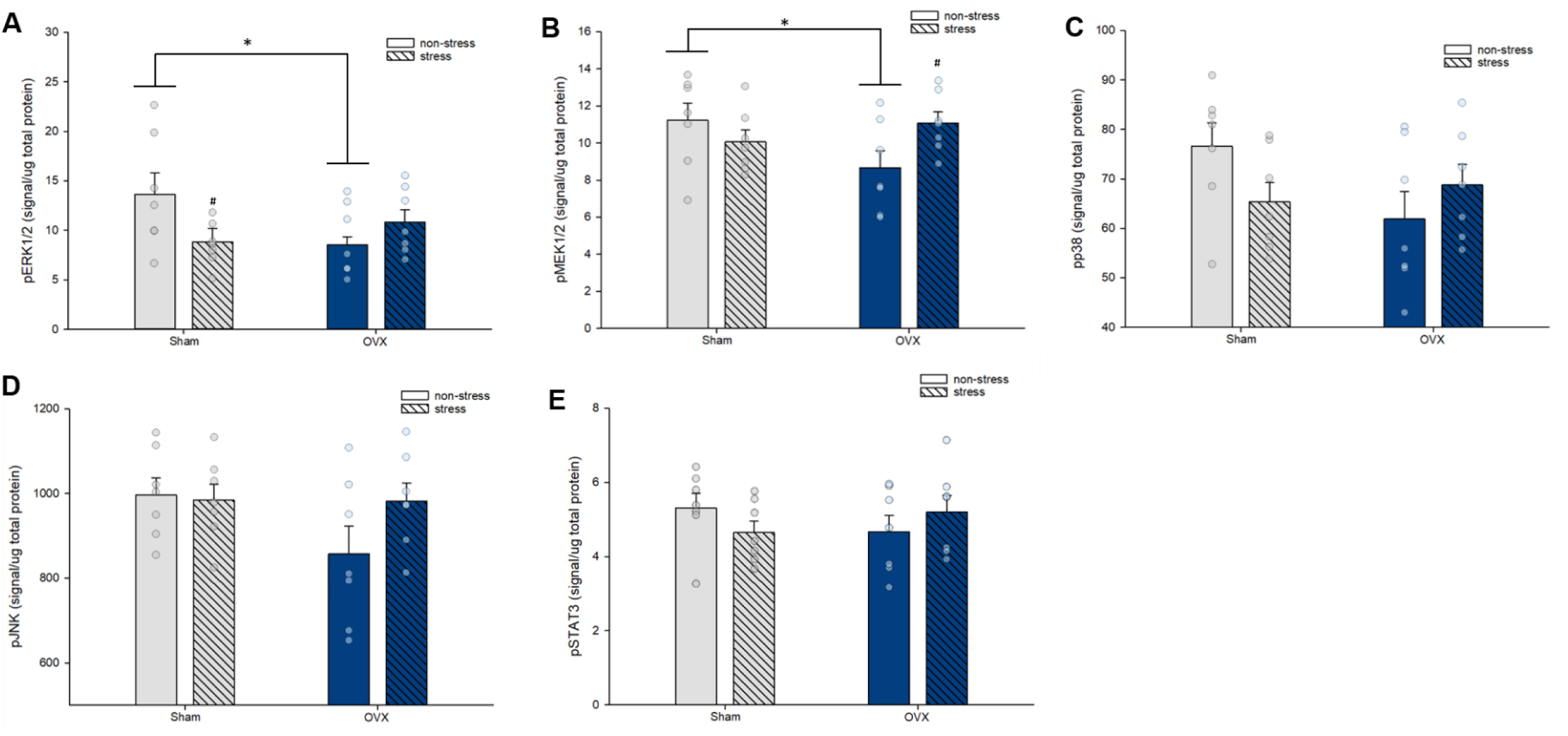
Cell signaling phosphoprotein expression in the frontal cortex, normalized by total protein levels. **(A)** Under non-stress conditions, ovariectomy decreased pERK1/2 expression, and stress exposure decreased pERK1/2 expression in sham-operated mice only; * indicates p = 0.032, significant difference between non-stress groups; # indicates p = 0.023, relative to non-stress sham-operated mice. **(B)** Under non-stress conditions, ovariectomy decreased pMEK1/2 expression, and stress exposure increased pMEK1/2 expression in ovariectomized mice only; * indicates p =0.032, significant difference between non-stress groups; # indicates p =0.042, relative to non-stress ovariectomized mice. There were no significant group differences in pp38 **(C),** pJNK **(D)**, and pSTAT3 **(F)** expression. Data in means + standard error of the mean. OVX, ovariectomized; ERK, extracellular signal-regulated kinase; JNK, c-Jun NH_2_-terminal kinase. n = 7/group. Data in means + standard error of the mean.

#### PSD-95 expression was not significantly affected by ovariectomy or sub-chronic stress exposure

Expression of the postsynaptic scaffolding protein PSD-95 was utilized as a synaptic marker and a proxy measure of excitatory synapse density (Wainwright et al., 2016). PSD-95 expression was previously found to be reduced by approximately 40% in the PFC of individuals with depression (Feyissa et al., 2009), Here, PSD-95 expression in the frontal cortex was not significantly affected by stress condition, ovarian status, nor their interaction (all p’s >0.4; **Table 5**).

**Table 5.** Postsynaptic density protein-95 expression in the prefrontal cortex.

| Group | PSD-95 (signal/ug protein) |
| --- | --- |
| Sham, non-stress | 376.11 ± 53.53 |
| Sham, stress | 412.56 ± 54.68 |
| OVX, non-stress | 415.88 ± 31.09 |
| OVX, stress | 445.36 ± 18.38 |

Data in means ± standard error of the mean. $n = 7/\text{group}$ . OVX, ovariectomized; PSD-95, postsynaptic density protein-95.

**Table 6.** Correlations between cell signaling phosphoproteins and cytokine concentrations in the frontal cortex.

|  | pERK1/2 | pMEK1/2 | pp38 | pJNK | pSTAT3 |
| --- | --- | --- | --- | --- | --- |
| **Frontal cortex** |  |  |  |  |  |
| **IL-1β** | -0.30<br>p=0.12 | 0.16<br>p=0.40 | -0.09<br>p=0.63 | 0.25<br>p=0.192 | -0.09<br>p=0.65 |
| **IL-2** | 0.51<br>p=0.005 | 0.69*<br>p<0.001 | 0.79*<br>p<0.001 | 0.57<br>p=0.002 | 0.71*<br>p<0.001 |
| **IL-4** | 0.17<br>p=0.40 | 0.08<br>p=0.67 | 0.25<br>p=0.19 | 0.10<br>p=0.60 | 0.11<br>p=0.59 |
| **IL-5** | 0.39<br>p=0.04 | 0.49<br>p=0.008 | 0.67*<br>p<0.001 | 0.40<br>p=0.034 | 0.65*<br>p<0.001 |
| **IL-6** | 0.20<br>p=0.30 | 0.53<br>p=0.004 | 0.46<br>p=0.014 | 0.49<br>p=0.009 | 0.46<br>p=0.015 |
| **IL-10** | 0.45<br>p=0.021 | 0.64*<br>p<0.001 | 0.69*<br>p<0.001 | 0.48<br>p=0.013 | 0.56<br>p=0.003 |
| **IL-12p70** | 0.55<br>p=0.002 | 0.65*<br>p<0.001 | 0.81*<br>p<0.001 | 0.489<br>p=0.008 | 0.68*<br>p<0.001 |
| **TNF-α** | 0.09<br>p=0.67 | 0.49<br>p=0.008 | 0.45<br>p=0.017 | 0.54<br>p=0.003 | 0.30<br>p=0.12 |
| **IFN-γ** | 0.43<br>p=0.026 | 0.70*<br>p<0.001 | 0.72*<br>p<0.001 | 0.60*<br>p=0.001 | 0.61*<br>p=0.001 |
| **CXCL1** | 0.41<br>p=0.03 | 0.70*<br>p<0.001 | 0.73*<br>p<0.001 | 0.80*<br>p<0.001 | 0.58*<br>p=0.001 |

Pearson's correlation coefficients (r) and p values. \* and grey background indicate significant correlations after correcting for multiple comparisons. IL, interleukin; CXCL1, chemokine (C-X-C motif) ligand 1; TNF-α , tumor necrosis factor-α ; IFN-γ , Interferon-γ .

#### Reduced latency to immobility in the forced swim test was associated with increased IL-6:IL-10 in the frontal cortex and hippocampus

Pearson’s correlations were performed between behavioral measures and markers of neuroinflammation that were significantly affected by stress exposure in an ovarian status-dependent manner. We found that reduced latency to immobility in the forced swim test was significantly associated with increased IL-6:IL-10 in the frontal cortex (r = - 0.43, p = 0.048; **Fig. 10A**) and hippocampus (r = - 0.43, p = 0.044; **Fig. 10B**). However, there were no significant correlations between latency to immobility and IL-1β concentrations in the frontal cortex (r = - 0.11, p = 0.6) or hippocampus (r = 0.04, p = 0.85).

**Figure 10.**
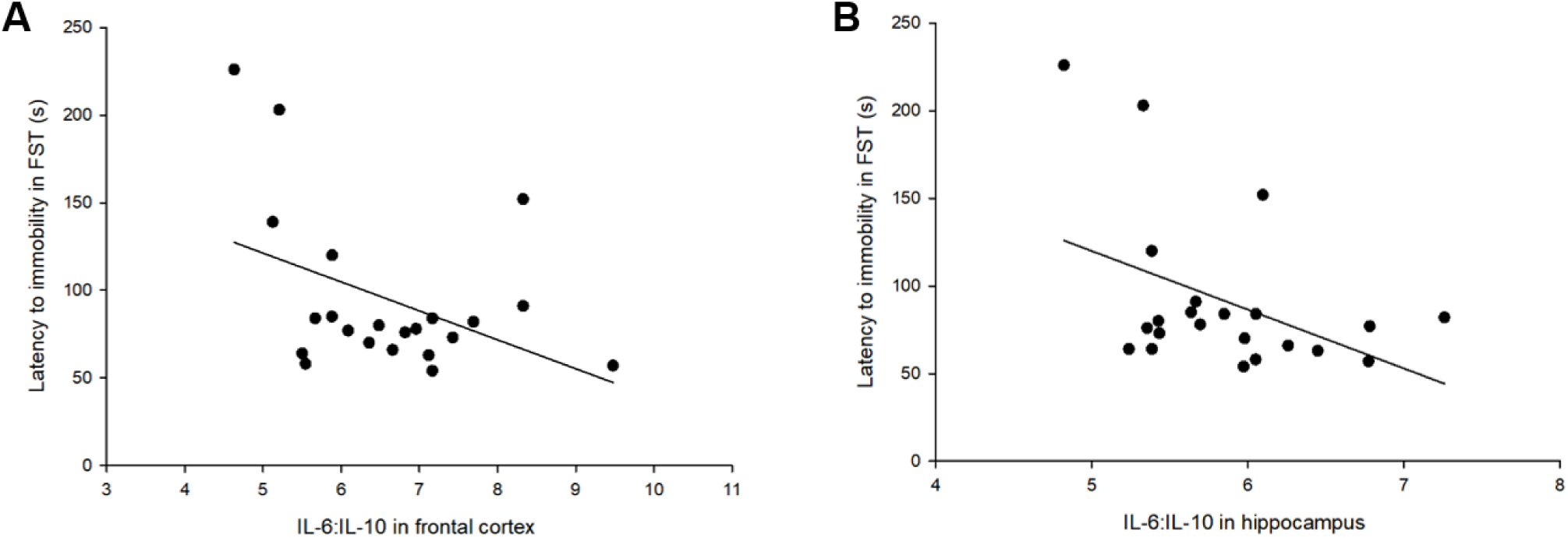
Association between passive-coping behavior and pro- to anti-inflammatory cytokine balance in the frontal cortex **(A)** and hippocampus **(B)**. Reduced latency to immobility was significantly associated with increased IL-6:IL-10 in the frontal cortex (**A**; r = - 0.43, p = 0.048) and hippocampus (**B**; r = - 0.43, p = 0.044). FST, forced swim test; IL, interleukin.

## Discussion

Here, we report profound effects of long-term ovariectomy on the central cytokine milieu, evidenced by reduced cytokine concentrations in the frontal cortex but increased cytokine concentrations in the hippocampus, in middle-aged female mice. Long-term ovariectomy also reduced phosphorylated ERK1/2 and MEK1/2 expression in the frontal cortex in middle age. Along with these neural changes in the frontal cortex and hippocampus, long-term ovariectomy resulted in a modest increase in depressive-like behavior under non-stress conditions in middle age. We further show ovarian status-dependent effects of sub-chronic stress exposure on cytokine concentrations in the frontal cortex and hippocampus, as seen by increased IL-1β and a shift toward a pro-inflammatory cytokine bias (IL-6:IL-10) in sham-operated mice only. Sub-chronic stress exposure also decreased expression of phosphorylated ERK1/2 in the frontal cortex in sham-operated mice only. Importantly, this was coupled with a greater behavioral susceptibility to sub-chronic stress in sham-operated mice. These data suggest that ovarian hormones exert a powerful influence on the neuroimmune environment in a region-dependent manner, and may dictate certain behavioral and neuroinflammatory consequences of sub-chronic stress exposure.

### Long-term ovariectomy modifies the cytokine signature under non-stress conditions and in response to sub-chronic stress exposure

We observed brain region-specific effects of long-term ovariectomy (9 months) on cytokine concentrations, with reductions in the frontal cortex and modest elevations in the hippocampus. Indeed, more than 50% of the variance within the frontal cortex cytokine data was explained by Principal Component 1 (PC1), which appeared to be largely accounted for by ovarian status. Ovarian status also affected the hippocampal cytokine milieu, although to a lesser extent than in the frontal cortex. Overall, the principal component loadings indicated 7 overlapping cytokines were important contributors to the first principle component (PC1) that was modified by long-term ovariectomy albeit in opposing directions between the frontal cortex and hippocampus (IL-2, IL-4, IL-5, IL6, IL-10, IL-12p70, TNF-α). In terms of individual ANOVA analyses, more distinct sets of cytokines were affected by long-term ovariectomy in the two brain regions, with the exception of IL-6 which was increased in the hippocampus but decreased in the frontal cortex. The effects of sub-chronic stress exposure on cytokine concentrations in the frontal cortex and hippocampus were less pronounced. This is not surprising given the sub-chronic nature of the stress paradigm (6 days), thus we would expect longer durations of stress exposure to result in larger alterations in the cytokine milieu. However, component loadings indicate IL-1β, TNF-α and the chemokine CXCL1 as important contributors to PC2 in the hippocampus, which showed significant effects of sub-chronic stress exposure. Overall, we do not observe a clear division of pro-inflammatory or anti-inflammatory cytokines, nor of T helper (Th) 1 and Th2 type cytokines, in the contribution to PCs. This may not be surprising, as although cytokine responses can be biphasic (i.e. an initial pro-inflammatory response followed by an anti-inflammatory response), simultaneous pro- and anti-inflammatory responses can also be observed. Indeed, a meta-analysis indicated that individuals with depression display increased concentrations of IL-10, an anti-inflammatory Th2 type cytokine, in addition to increased pro-inflammatory cytokines (Köhler et al., 2017), perhaps representing a compensatory effect. It is also important to note that all mice underwent behavioral testing, including the forced swim test which is an acute stressor. Therefore, it is plausible that exposing non-stress controls to behavioral testing may have masked some of the effects of sub-chronic stress exposure on cytokine concentrations.

An intriguing finding was that ovarian status impacted the neuroinflammatory consequences of sub-chronic stress exposure, with overall more pronounced effects in sham-operated mice. Specifically, in the frontal cortex and hippocampus of sham-operated mice only, stress exposure increased concentrations of the pro-inflammatory IL-1β and resulted in a shift toward a pro-inflammatory cytokine bias (IL-6:IL-10). The exaggerated neuroinflammatory response to stress in sham-operated mice was observed in tandem with a greater behavioral susceptibility to sub-chronic stress exposure (discussed below). Our findings in sham-operated females corroborate studies in male subjects, in which IL-1β was increased by chronic stress exposure in the brain, and further found to be directly implicated in the behavioral consequences of stress exposure (Goshen et al., 2008; Goshen and Yirmiya, 2009; Wohleb et al., 2014). Further, the role of IL-6 in depression and stress-based models is well-established (reviewed in Hodes et al., 2016), and a shift towards a higher IL-6: IL-10 ratio is associated with passive-coping in response to social defeat stress in male rats (Wood et al., 2015). Thus, our findings suggest that in middle-aged females, ovarian hormones may potentiate the inflammatory consequences of stress exposure. Given that ovariectomy was performed in young-adulthood, sham-operated groups more closely approximate what occurs in normal transition to reproductive senescence. Thus, the greater susceptibility to sub-chronic stress exposure in sham-operated mice may be linked to the endogenous changes in ovarian hormones at this time. Indeed, this mirrors the human condition in which the transition to menopause is associated with increased risk for depression (Cohen et al., 2006; Freeman et al., 2004).

Ovarian hormones, especially 17β-estradiol but also progesterone, have well established anti-inflammatory properties (Bruce-Keller et al., 2000; He et al., 2004; Lei et al., 2014; Vegeto et al., 2008). Consistent with this are the observed effects of long-term ovariectomy to increase IL-6:IL-10 ratio in the frontal cortex and IL-6 concentrations in the hippocampus under non-stress conditions. Given the role of IL-6 in MDD (Dowlati et al., 2010; Liu et al., 2012), these observations may be related to the modest increase in depressive-like behavior induced by long-term ovariectomy under non-stress conditions, which is in line with findings in humans (Schmidt et al., 2015; Wilson et al., 2018). However, in the frontal cortex we also found that ovariectomy reduced concentrations of other proinflammatory cytokines including IFN-γ and TNF-α. Further, in both the frontal cortex and hippocampus we observed an exaggerated neuroinflammatory response to stress in sham-operated, rather than ovariectomized mice, highlighted by increased IL-1β and increased IL-6:IL-10 ratio. Importantly, the anti-inflammatory and neuroprotective effects of estrogens have been investigated mostly in younger females and in models of brain injury or stroke, and in response to acute inflammatory challenges or *in vitro* (Selvamani and Sohrabji, 2010; Suzuki et al., 2009; Vegeto et al., 2003). Age is a critical consideration here, as studies using older females have observed neurotoxic effects of estrogens in similar models (Johnson and Sohrabji, 2005; Nordell et al., 2003; Selvamani and Sohrabji, 2010), partially consistent with our finding here that exposure to sub-chronic stress resulted in a larger neuroinflammatory response in sham-operated middle-aged mice. Future studies need to consider age as a variable and should introduce 17β-estradiol and/or progesterone replacement to further clarify the role of ovarian hormones in the neuroinflammatory outcomes of stress exposure. Additionally, it would be important that future studies investigate whether changes in estrous cyclicity in middle-aged animals can affect the outcomes of stress exposure, as the current study was not powered to directly answer this question.

### Long-term ovariectomy and sub-chronic stress exposure interact to affect ERK signaling

Long-term ovariectomy reduced phosphorylated ERK1/2 and its upstream activator MEK1/2 in the frontal cortex under non-stress conditions, and stress exposure reduced pERK1/2 expression in sham-operated mice only. The observed reduction with long-term ovariectomy may not be surprising, as this group had significantly lower circulating concentrations of estradiol, which can rapidly activate ERK1/2 (Bi et al., 2002; Fernandez et al., 2008). Perhaps more intriguing than baseline differences in pERK1/2 expression were the differential effects of stress exposure, as abnormal ERK signaling has been observed in post-mortem brains of individuals with MDD that died of suicide (Duric et al., 2010; Dwivedi et al., 2006, 2001; Hsiung et al., 2003; Labonté et al., 2017). Notably, in human studies there are reports of reduced ERK1/2 expression and activation in the prefrontal cortex and hippocampus of males and females with MDD (Dwivedi et al., 2006, 2001; Hsiung et al., 2003), although data was not stratified by sex. Our results contrast another study in which chronic stress exposure (21 days) increased pERK1/2 in the ventromedial PFC of female mice (Labonté et al., 2017), but this study used young adult females whereas our subjects were middle-aged. Collectively, past studies indicate that dysregulated ERK signaling is implicated in depression and the outcome of stress exposure and our current findings extend previous work to suggest that ovarian hormones can influence the outcomes of stress exposure on ERK signaling in females. Moreover, we observe significant positive correlations between several cytokines and cell signaling phosphoproteins in the frontal cortex. This is to be expected as not only do cytokines activate all pathways examined in this study, but MAPKs are also important regulators of cytokine production (Cargnello and Roux, 2011; Johnson and Lapadat, 2002). Using specific MAPK inhibitors, future experiments could assess the directionality and causality of the observed relationships between cytokines and cell signaling phosphoproteins. In the present study, due to limited tissue, we only examined cell signaling phosphoproteins in the frontal cortex, therefore future studies should examine these signaling pathways in the hippocampus.

Long-term ovariectomy and stress did not significantly affect PSD-95 expression in the frontal cortex in the present study. This finding is consistent with previous studies indicating that although PSD-95 expression is sensitive to transient fluctuations in ovarian hormones (Spencer et al., 2010), it is not modified two weeks post ovariectomy (Spencer-Segal et al., 2012; Waters et al., 2015, 2009; Zhang et al., 2010). Several studies have found reductions in PSD-95 expression in response to chronic stress exposure (Kallarackal et al., 2013; Kim and Leem, 2016; Pacheco et al., 2017) suggesting that the shorter duration of stress exposure used in this study may not be sufficient to produce significant modifications to PSD-95 expression in the frontal cortex. Future studies should also examine the effects on PSD-95 expression in the hippocampus and with longer durations of stress exposure.

### Ovarian status influenced behavior under non-stress conditions and in response to sub-chronic stress exposure

Under non-stress conditions, long-term ovariectomy alone impacted some measures of depressive-like behavior, as seen by reduced latency to immobility in the forced swim test and a trend toward reduced sucrose consumption. These behavioral differences may be linked to the reported effects of long-term ovariectomy on cytokine concentrations in the brain. However, the depressogenic effects of long-term ovariectomy were overall limited, as we did not observe significant differences between sham-operated and ovariectomized mice in any other behavioral measure. This is partially consistent with other studies in which the anxiogenic and depressogenic effects of ovariectomy (Estrada-Camarena et al., 2011; Li et al., 2014) can recover after more prolong periods of ovarian hormone deprivation (de Chaves et al., 2009; Estrada-Camarena et al., 2011).

Exposure to sub-chronic stress reduced latency to immobility in the forced swim test in sham-operated mice only, pointing to a greater susceptibility to stress-induced passive-coping behavior. However, this effect was not seen across all measures, as sub-chronic stress exposure increased anhedonia-like behavior and reduced body mass regardless of ovarian status, suggesting similar susceptibilities in these domains. Further, we do not observe significant group differences in the percentage of time spent immobile in the forced swim test, suggesting overall modest behavioral effects of stress exposure, likely due to the sub-chronic nature of the paradigm (6 days). In the forced swim tests, it is also plausible that ovariectomized mice had reached a floor effect under non-stress conditions, leaving no room for sub-chronic stress exposure to further reduce latency to immobility. However, the effect of stress to reduce latency to immobility in sham-operated mice was observed in concert with an exaggerated neuroinflammatory response to stress and a reduction in pERK1/2 expression. Therefore, in comparison with ovariectomized mice, we observe more robust effects of sub-chronic stress exposure in sham-operated mice across several measures, perhaps supporting the interpretation of greater overall susceptibility to the effects of sub-chronic stress exposure. Indeed, reduced latency to immobility was significantly correlated with increased IL-6:IL-10 ratio in the frontal cortex and hippocampus, suggesting that the greater susceptibility to stress-induced passive-coping behavior in sham-operated mice may be directly linked to the exaggerated neuroinflammatory response to stress. There are several possible mechanisms by which inflammation can ultimately influence depressive-like behavior. Notably, pro-inflammatory cytokines can affect multiple neurotransmitter systems, including serotonergic, glutamatergic, and dopaminergic systems (Felger, 2017; Haroon and Miller, 2017; Miller et al., 2014). For example, in male mice, IL-1β has been shown to decrease synaptic availability of serotonin by enhancing serotonin transporter activity, an effect that was directly linked to depressive-like behavior (Zhu et al., 2010). Further, adult hippocampal neurogenesis, which is implicated in stress resilience (Anacker et al., 2018), is reduced by inflammation (Ekdahl et al., 2003; Monje et al., 2003), and regulated directly by IL-1β (Zunszain et al., 2012). Future studies should investigate whether these mechanisms are implicated in the effects of ovarian hormones to influence stress-induced inflammation and behavior.

Our current findings agree with a previous study in young adult female mice in which exposure to a similar sub-chronic stress paradigm increased passive-coping behavior in sham-operated, but not ovariectomized mice (LaPlant et al., 2009). In contrast, other studies in rats and mice found that long-term ovariectomy increased depressive-like behavior after chronic stress exposure (Lagunas et al., 2010; Mahmoud et al., 2016). These dissimilarities may be reconciled by comparing the duration of exposure to stressors, which spanned 4-6 weeks in studies showing increased sensitivity to depressive-like behavior after long-term ovariectomy (Lagunas et al., 2010; Mahmoud et al., 2016), as opposed to 6 days in the current report and in LaPlant et al (2009), where increased sensitivity to stress is observed in sham-operated females. Thus, together with previous studies our current data indicate that ovarian hormones can confer either risk or resilience, depending on stress paradigm.

## Conclusions

In conclusion, long-term ovarian hormone deprivation significantly altered the neuroimmune environment in a brain region-specific manner, reduced expression of ERK pathway phosphoproteins in the frontal cortex, and modestly increased depressive-like behavior under non-stress conditions, in middle-aged mice. Importantly, ovarian status dictated some of the behavioral and neuroinflammatory outcomes of sub-chronic stress exposure. Specifically, sham-operated mice had a greater neuroinflammatory response to stress as seen by increased concentrations of IL-1β and increased IL-6:IL-10 in the frontal cortex and hippocampus. We also observed stress-induced reductions in pERK1/2 expression in the frontal cortex of sham-operated mice only. This was coupled with an enhanced behavioral susceptibility to sub-chronic stress exposure in sham-operated mice, as seen by increased passive-coping behavior in the forced swim test. It is important to note ovariectomy was performed in young adulthood and outcomes were examined in middle-age. Therefore, our sham-operated middle-aged mice, tested at a time of transition to reproductive senescence, are perhaps a closer model of the menopausal transition. With this in mind, it may not be surprising that the outcomes of stress exposure were more robust in sham-operated mice, as this mirrors the human condition where the transition to menopause carries an increased risk for depression. These findings underscore the importance of considering the immunomodulatory properties of ovarian hormones in stress and depression research.

